# Distinct roles for thymic stromal lymphopoietin (TSLP) and IL-33 in experimental eosinophilic esophagitis

**DOI:** 10.1101/2025.02.25.640192

**Authors:** Anish Dsilva, Ariel Wagner, Michal Itan, Natalie Rhone, Shmulik Avlas, Yaara Gordon, Natalie Davidian, Shraddha Sharma, Elizaveta Razravina, Israel Zan-Bar, Jane R. Parnes, Kevin S. Gorski, Joseph D. Sherrill, Chen Varol, Steven F. Ziegler, Marc E. Rothenberg, Ariel Munitz

**Affiliations:** Department of Clinical Microbiology and Immunology, Faculty of Medical and Health Sciences, Tel-Aviv University, Israel; Early Development, Amgen, Thousand Oaks, California, USA; Clinical Biomarkers and Diagnostics, Amgen, South San Francisco, California, USA; Translational Science and Experimental Medicine, Research & Early Development, Respiratory & Immunology, BioPharmaceuticals R&D, AstraZeneca, Gaithersburg, Maryland, USA; Research Center for Digestive Tract and Liver Diseases, Sourasky Medical Center, Tel Aviv, Israel; Center for Fundamental Immunology, Benaroya Research Institute, Seattle, WA, USA; Division of Allergy/Immunology, Department of Pediatrics, Cincinnati Children’s Hospital Medical Center and the University of Cincinnati College of Medicine, Cincinnati, Ohio, USA

**Author notes:** **Corresponding author:** Ariel Munitz, PhD, Department of Clinical Microbiology and Immunology, The Sackler School of Medicine, Tel Aviv University, Ramat Aviv 69978, Israel. **Grant Support:** Ariel Munitz is supported by the US-Israel Bi-national Science Foundation (grant no. 2023029 with MER), by the Israel Science Foundation (grants no. 328/24 and 542/20), the Israel Cancer Research Fund, the Israel Cancer Association Avraham Rotstein Donation, the Cancer Biology Research Center (TAU), the Azrieli Foundation Canada-Israel and External sponsored scientific research grants from AstraZeneca. Ariel Munitz and Chen Varol are supported by the Azrieli Foundation Canada-Israel. Marc E. Rothenberg is supported by grants [R37 AI045898, R01 AI124355, U19 AI070235, and P30 DK078392 (Gene and Protein Expression Core); the Campaign Urging Research for Eosinophilic Disease (CURED); the Sunshine Charitable Foundation and its supporters, Denise A. Bunning and David G. Bunning]. **Disclosures:** Ariel Munitz is a consultant for Glaxo Smith Kline, Astra Zeneca, Sanofi, Oravax, Sartorious and is an inventor of patents owned by the Tel Aviv University. Marc E. Rothenberg is a consultant for Pulm One, Spoon Guru, ClostraBio, Serpin Pharm, Celldex, Uniquity Bio, EnZen Therapeutics, Bristol Myers Squibb, Astra Zeneca, Pfizer, Glaxo Smith Kline, and Regeneron/Sanofi, and Guidepoint and has an equity interest in the first seven listed plus in Allakos and Santa Ana Bio, and royalties from reslizumab (Teva Pharmaceuticals), PEESSv2 (Mapi Research Trust), and UpToDate. M.E.R. is an inventor of patents owned by Cincinnati Children’s Hospital Medical Center. The remaining authors disclose no conflicts. Jane R. Parnes and Kevin S. Gorski are employees of Amgen and may own stock or stock options in Amgen. Joseph D. Sherrill is an employee of AstraZeneca and may own stock or stock options in AstraZeneca.

**Keywords:** Allergy, Eosinophilic Esophagitis, TSLP, IL-33

## Abstract

**Rationale:** Thymic stromal lymphopoietin (TSLP) and IL-33 are alarmins implicated in EoE pathogenesis by activating multiple cells including mast cells (MCs). Whether TSLP or IL-33 have a role in EoE and whether their activities are distinct requires further investigation.

**Methods:** Experimental EoE was induced in wild type (WT) *Il33^-/-^*and *Crlf2^-/-^* mice. TSLP or IL-5 were neutralized using antibodies. Esophageal histopathology was determined by H&E, anti-Ki67, anti-CD31 and anti-MBP staining. Esophageal RNA was subjected to RNA sequencing. Bone marrow-derived MCs were activated with TSLP and IL-13 was determined (ELISA)

**Results:** TSLP and IL-33 were overexpressed in human and experimental EoE. Human and mouse esophageal MCs displayed the highest level of *Crlf2* (TSLPR) compared to other immune cells. *Crlf2^-/-^* mice were nearly-completely protected from EoE, and TSLP neutralization resulted in decreased basal cell proliferation, eosinophilia, lamina propria thickening and vascularization. Induction of experimental EoE in *Il33^-/-^* mice resulted in reduced eosinophilia but no alterations in tissue remodeling were observed compared to WT mice. RNA sequencing revealed that TSLP regulates the expression of key genes associated with human EoE (e.g. eotaxins*, Il19, Klk5, Flg, Il36rn, Il1r2*) and suggest a role for TSLP in regulating IL-1 signaling, barrier integrity and epithelial cell differentiation. Experimental EoE was characterized by a MC-associated gene signature and elevated MCs. Activation of MCs with TSLP resulted in secretion of IL-13.

**Conclusion:** TSLP and IL-33 have non-redundant functions in experimental EoE. This study highlights TSLP as an upstream regulator of IL-13 and a potential therapeutic target for EoE.

## Introduction

EoE is a chronic, food antigen-induced T helper type 2 (Th2)-associated, allergic- and immune-mediated disease. It is characterized histologically by eosinophil-predominant mucosal inflammation and clinically by esophageal dysfunction^1^. Emerging evidence places the esophageal epithelium at the center of EoE pathogenesis^2^. Esophageal epithelial cells are a source of the thymic stromal lymphopoietin (TSLP) and interleukin (IL)-33, key alarmin cytokines that are responsible for allergic sensitization and upstream regulators of Th2 cytokines including IL-13^2^. IL-13 is a critical effector cytokine in EoE that stimulates loss of epithelial cell differentiation and increased proliferation^3^. Esophageal mast cells (MCs), as well as a subset T cells, are a prominent source of IL-13 in EoE^4,5^. Thus, defining the role of molecular and cellular pathways upstream of IL-13-mediated pathology in EoE is timely and important since such pathways can serve as new therapeutic targets.

TSLP and IL-33 have been implicated in various allergic diseases including EoE. Nonetheless, they mediate their activities via distinct mechanisms. TSLP is a pleiotropic cytokine that signals via a high-affinity TSLP receptor (R) complex, which is a heterodimer comprising cytokine receptor-like factor 2 (CRLF2/TSLPR) and the IL-7R α-chain^6^. Genome-wide association studies (GWAS) have repeatedly determined that genetic susceptibility for EoE is linked to genetic variants in the *TSLP* gene at 5q22^7^. TSLP protein is broadly expressed throughout the esophageal epithelium^8,9^. On the transcriptome level, high *Tslp* expression characterizes a subgroup of patients with EoE that represents the most inflammatory type 2-associated subgroup of EoE^10^. Mechanistically, TSLP directly upregulates co-stimulatory molecules on dendritic cells, which are required for the polarization of T cells into Th2 cells^11,12^. Furthermore, TSLP can increase the proliferation of Th2 cells and enhance the release of Th2 cell-associated cytokines particularly in the context of EoE^13^, as well as release of chemokines from MCs, innate lymphoid cells (ILCs), and macrophages^13–17^. TSLP has been also suggested to activate basophils to promote EoE, and TSLP gain-of-function polymorphisms were associated with increased basophil responses in patients with EoE^18^.

IL-33 is a member of the IL-1 cytokine family^19^ and is constitutively present in the nuclei of distinct populations of epithelial and endothelial cells. The receptor for IL-33 consists of IL1RL1/ST2 (termed herein ST2), which is specific to IL-33, and a shared signaling chain with the IL-1 family, IL-1RAcP^20^. While the role of IL-33 in EoE is largely unknown, accumulating evidence suggests pathological roles for this cytokine. Increased expression of IL-33 mRNA was found in the esophageal mucosa of EoE patients and IL-33 protein was detected in epithelial cells, MCs, and additional CD45^-^ cells^21,22^. Furthermore, expression of IL-33 is induced in undifferentiated, non-dividing esophageal epithelial cells in EoE^21^, and was shown to be activated via intracellular cleavage by ripoptosome activation following epithelial cell damage^23^. Recently, transgenic overexpression of IL-33 in esophageal epithelial cells is sufficient to induce an EoE-like disease in mice^24^ and intraperitoneal injection of IL-33 induced IL-13-dependent esophageal eosinophilia and epithelial cell proliferation^22^.

Herein, we aimed to define the potential roles of TSLP or IL-33 in experimental EoE, and whether their activities are distinct. Using an IL-13-dependent mouse model of experimental EoE that mimics the histopathological and molecular characteristics of human EoE^25^, we examined the roles of TSLP and IL-33 in disease progression. Our findings reveal non-redundant functions for these cytokines in driving the pathological features of EoE. TSLP regulated esophageal basal cell hyperproliferation, eosinophilia, lamina propria thickening and vascularization whereas IL-33 regulated only esophageal eosinophilia only. RNA sequencing analyses revealed that TSLP regulated key genes associated with IL-13-signaling and human EoE (e.g. eotaxins*, Il19, Klk5, Flg, Il36rn, Il1r2*) and that TSLP promoted IL-13 secretion from mucosal MCs. These data suggest a distinct and broad role for TSLP in experimental EoE that may be mediated by MCs and IL-13.

## Materials and Methods

### Experimental EoE

Mice (male c57BL6 mice, 6-8 weeks) were skin sensitized (15μl 1% OXA, Sigma #E0753, dissolved in acetone) on both ear flanks (60 μl per mouse). After six additional skin challenges (0.5% OXA in acetone), the mice were bled and serum IgE was quantitated (ELISA, BD Bioscience, USA). The mice were intraesophageally challenged (day 18, 200µl OXA, 1% in a 1:2 ratio of olive oil and 95% alcohol, respectively) using a plastic feeding tube that reaches the mid esophagus (INSTECH #ITH-FTP-22-25-50, 22gaX25mm) and has been modified for intraesophageal administration by puncturing 8 holes with a 14G needle^25^. On day 36, the mice were euthanized. A thorough explanation of the methods used in this study and analyses can be found in the supplemental file online.

## Results

### Increased TSLP and IL-33 expression in experimental EoE

IL-33 and TSLP have been implicated in the pathophysiology of EoE^8,13,26^. Thus, we were interested to determine whether levels of IL-33 and TSLP are increased in experimental EoE (Figure 1A). Intra-esophageal challenge of oxazolone to skin sensitized mice resulted in rapid and robust elevation of IL-33 in the esophagus, which was evident as early as day 19 (2 days post-intraesophgeal challenge) and remained elevated throughout the experiential protocol (Figure 1B). In contrast, TSLP expression was increased at later stages following intraesophageal challenge peaking at day 29 (Figure 1B). The kinetics of IL-33 and TSLP expression following exposure to oxazolone was different in the esophagus compared to the skin. Skin sensitization induced TSLP (but not IL-33) expression, which declined with the cessation of exposure to oxazolone (Figure 1C). IL-33 levels decreased initially following skin sensitization but returned to baseline levels during the period of intraesophageal challenges (days 17-29, Figure 1C).

**Figure 1.**
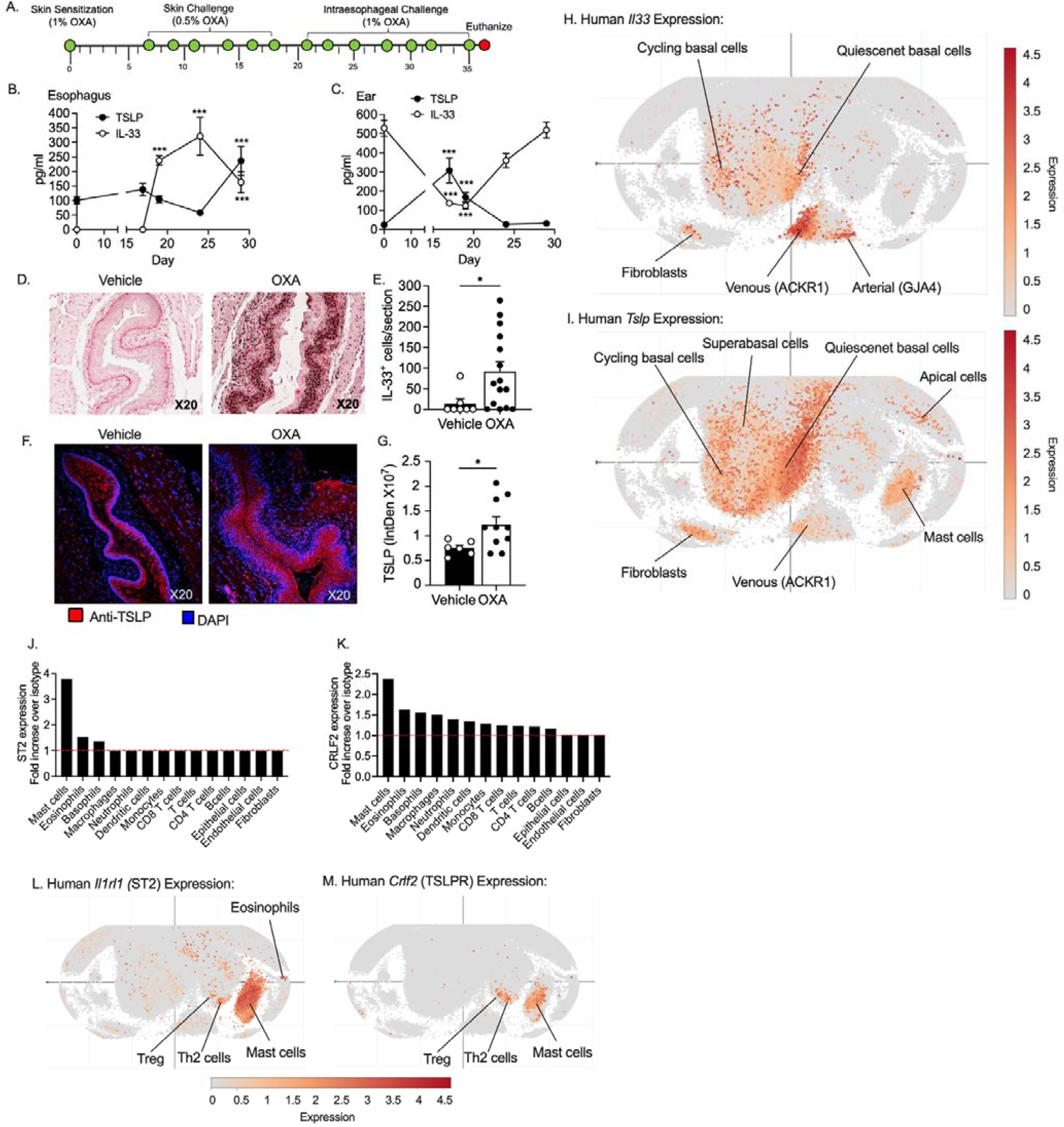
Expression of TSLP, IL-33 and their receptors in experimental and human EoE. Schematic representation of the experimental EoE model (A). Kinetic analysis of TSLP (black circle) and IL-33 (white circle) expression in the esophagus (B) and ear (C) throughout the experimental EoE protocol. Esophageal samples were obtained and the expression of IL-33 and TSLP were determined at day 36 by immunohistochemistry or immunofluorescence, respectively (D, F). Thereafter, the number of IL-33^+^ cells or the density of TSLP staining in the esophagus were quantified (E, G). Analysis of IL-33 and TSLP expression in human EoE samples was obtained from the human single cell portal (H, I). The expression of mouse IL1RL1 (ST2) and CRLF2 was determined in single cell suspensions obtained from the esophagus (J, K) using polychromatic flow cytometry. Data are shown as fold increase of anti-ST2 (J) or anti-CRLF2 (K) over isotype control. The red line represents no fold induction when calculating anti-ST2 or anti-CRLF2 over isotype control. Analysis of *Il1rl1* and *Crlf2* expression in human EoE samples was obtained from the human single cell portal (L, M). Cells in the portal were annotated using the single cell portal (https://singlecell.broadinstitute.org/single_cell). In B, C, data are presented as mean ± SEM from n= 6-8 mice per time point. In E, G, each dot represents one mouse. In H, I, L and M each dot represents one cell where the intensity of the color represents the expression of *Il33* or *Tslp*. *\*-p<0.05, **-p<0.01, ***-p<0.001*.

Immunohistochemical and immunofluorescence staining for IL-33 and TSLP, respectively, in the esophagus demonstrated that IL-33 was nearly absent under baseline conditions but highly upregulated, predominantly in esophageal epithelial cells, following intraesophageal challenge (Figure 1D-E). TSLP was expressed under baseline conditions in the esophagus and increased by day 36 in epithelial and non-epithelial cells following intraesophageal challenge (Figure 1F-G).

To translate these findings into human disease, the expression of IL-33 and TSLP were examined in a publicly available database of single cell RNA sequencing (scRNAseq) from esophageal biopsies of patients with EoE^27^. IL-33 was highly expressed by venous endothelial cells (ACKR1^+^), arterial cells (GJA4^+^) as well as quiescent and cycling basal cells and to lesser extent by fibroblasts (Figure 1H). TSLP displayed a broader expression pattern in the esophagus that included quiescent, cycling basal epithelial cells and suprabasal epithelial cells, as well as fibroblasts and mast cells (Figure 1I). Venous endothelial cells, apical cells and KLRG1^+^ CD8^+^ effector T cells also expressed TSLP to lesser extent (Figure 1I).

### Esophageal mast cells (MCs) highly express receptors for IL-33 and TSLP

To better understand which cells may respond to IL-33 and/or TSLP in the esophagus, cell surface expression of IL1RL1 (ST2) and cytokine receptor like factor 2 (CRLF2, which encodes for TSLPR) were determined in enzymatically digested mouse esophageal cell suspensions (See gating strategy in Figure S1A). ST2 and CRLF2 were expressed the highest in mast cells compared to other esophageal cells (Figure 1J-K and S1B-C). The expression of ST2 was also detected on eosinophils and basophils, whereas CRLF2 expression displayed a broader expression pattern and was identified in additional immune cells such as dendritic cells (DC), T cells and monocytes (Figure 1J-K).

Analysis of *Il1rl1* and *Crlf2* gene expression in human single-cell RNA sequencing data^27^, demonstrated that *Il1rl1* and *Crlf2* display a similar expression pattern to that observed in mouse cells as both genes were highly expressed in mast cells and to a lesser extent in Th2 and T regulatory cells (Figure 1L-M).

### IL-33 regulates the infiltration of eosinophils in experimental EoE

To examine the role of IL-33 in EoE, experimental EoE was induced in *Il33^-/-^* mice^28^. OXA-challenged *Il33^-/-^* mice displayed decreased infiltration of intraepithelial and lamina propria eosinophils in the esophagus (Figure 2A-B), Interestingly, basal cell proliferation (marked by Ki-67 staining, Figure 2C-D) and subsequent epithelial cell thickening were independent of IL-33 (Figure 2E-F). Furthermore, esophageal lamina propria thickness, (Figure 2G) and esophageal vascularization (determined by anti-CD31 staining, were independent of IL-33 (Figure 2H-I). These data suggest a limited role for IL-33 in experimental EoE.

**Figure 2.**
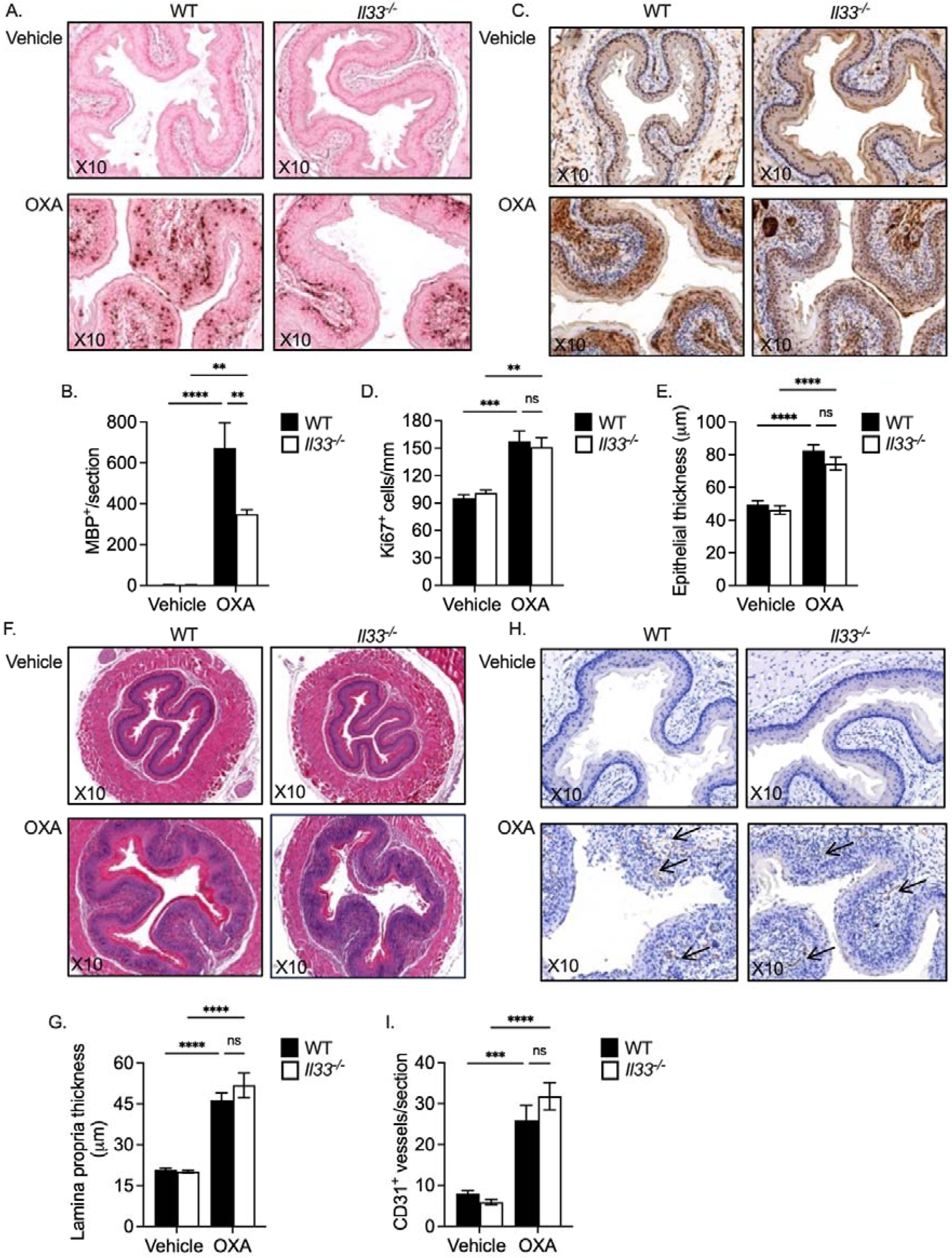
IL-33 regulates the accumulation of eosinophils in experimental EoE. Experimental EoE was induced in wild type (WT) and *Il33^-/-^* mice using oxazolone (OXA). On day 36, the mice were euthanized, and esophageal tissues were fixed, paraffin embedded, and slides were generated. The slides were stained with anti-major basic protein (MBP, A) and esophageal eosinophils were quantified (B). Epithelial cell proliferation was determined using anti-Ki67 staining (C, D). H&E-stained slides were analysed for lamina propria (E-F) and esophageal epithelial (G) thickness. Esophageal vascularization was determined using anti-CD31 staining (H, I). Representative photomicrographs of anti-MBP (A), anti-Ki-67 (C), H&E (F) and anti-CD31 (H) are shown. Data are presented as mean ± SEM and are representative of n=3 experiments conducted with 10-12 mice per group, ns-nonsignificant *\*\*-p<0.01, ***-p<0.001, ****-p<0.0001*.

### A key role for TSLP in experimental EoE

To examine the role of TSLP in EoE, experimental EoE was induced in *Crlf2^-/-^* mice^11^. OXA-challenged *Crlf2^-/-^* mice displayed no infiltration of eosinophils to the esophagus (Figure 3A-B), Furthermore, basal cell proliferation (marked by Ki-67 staining, Figure 3C-D) and subsequent epithelial cell thickening were completely dependent on TSLP (Figure 3E-F). Esophageal lamina propria thickness, (Figure 3G) and esophageal vascularization, were completely dependent of TSLP as well (Figure 3H-I). Importantly, increased expression of IL-33 following induction of experimental EoE was TSLP-dependent (Figure 3J). However, expression of TSLP was independent of IL-33 since oxazolone-challenged *Il33^-/-^*mice displayed elevated TSLP expression (Figure 3K-L).

**Figure 3.**
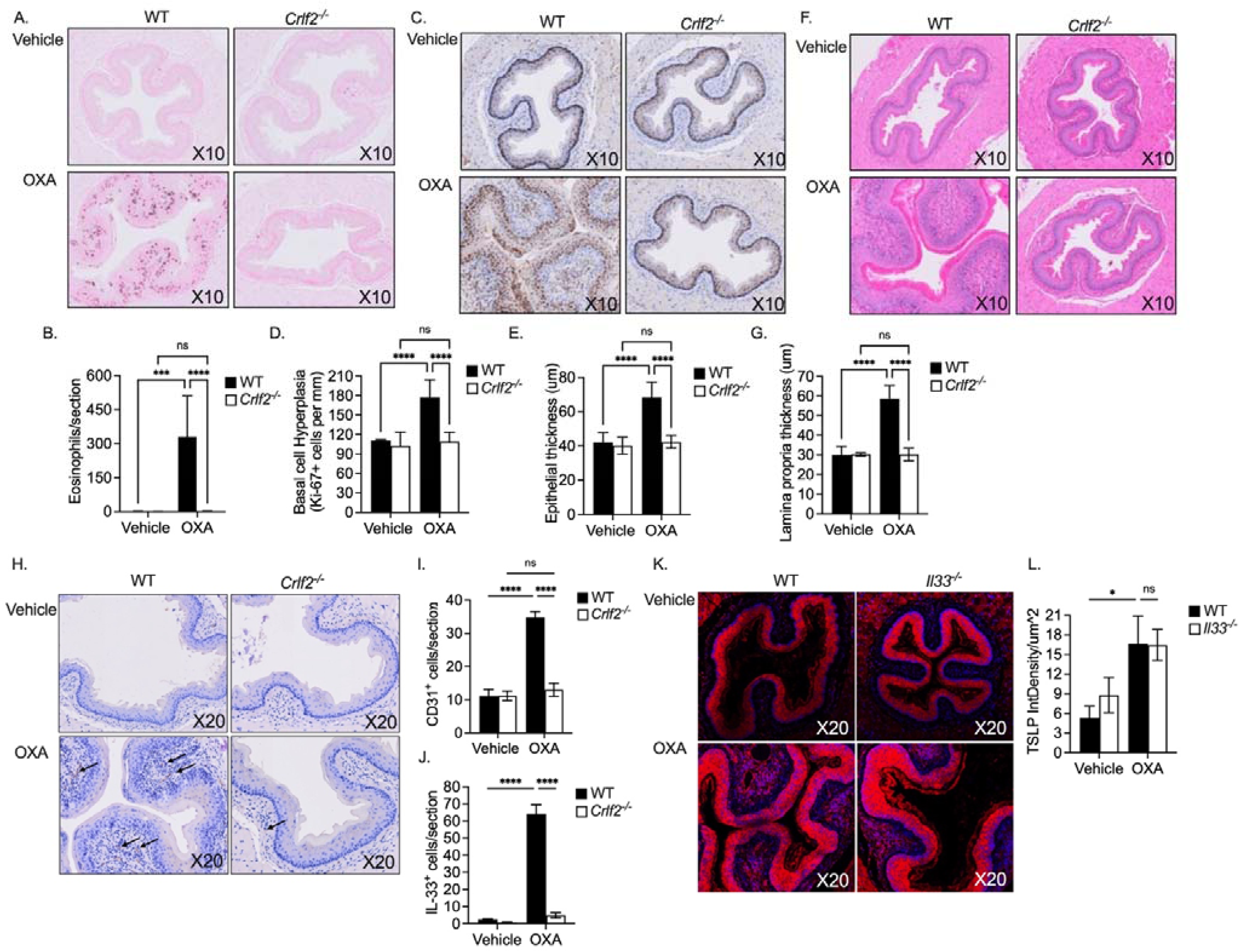
Experimental EoE is critically regulated by TSLP receptor. Experimental EoE was induced in wild type (WT) and *Crlf2^-/-^* mice using oxazolone (OXA). On day 36, the mice were euthanized, and esophageal tissues were fixed, paraffin embedded, and slides were generated. The slides were stained with anti-major basic protein (MBP, A) and esophageal eosinophils were quantified (B). Epithelial cell proliferation was determined using anti-Ki67 staining (C, D). H&E-stained slides were analysed for assessment of epithelial thickness (E-F) and lamina propria thickness (G). Esophageal vascularization was determined using anti-CD31 staining (H, I). Expression of IL-33^+^ cells was determined using anti-IL-33 immunohistochemistry (J). Experimental EoE was induced in wild type (WT) and *Il33^-/-^* mice using oxazolone (OXA). Expression of TSLP was determined using anti-TSLP by immunofluorescence (K-L). Representative photomicrographs of anti-MBP (A), anti-Ki-67 (C), H&E (F) and anti-CD31 (H) and anti-TSLP (K) are shown. Data are presented as mean ± SEM and are representative of n=3 experiments conducted with 10-12 mice per group, ns-nonsignificant *\*\*-p<0.01, ***-p<0.001, ****-p<0.0001*.

### Experimental EoE is independent of eosinophils

The absence of infiltrating eosinophils in the esophagus of oxazolone-challenged *Crlf2^-/-^* mice raised the possibility that decreased histopathology may be indirectly due to lack of eosinophils. To examine this possibility, eosinophils were depleted using anti-IL-5 antibodies (Figure S2). Anti-IL-5 treatment decreased esophageal eosinophil levels (Figure S2B). Despite this, oxazolone-challenged anti-IL-5-reated mice displayed similar levels of basal cell proliferation, epithelial thickening and edema (Figure S2C-E). Thus, experimental EoE is independent of eosinophils and decreased esophageal pathology in *Crlf2^-/-^* mice is eosinophil-independent.

### Pharmacological blockade of TSLP alleviates experimental EoE

TSLP can potentially regulate skin sensitization by governing B cell activation and subsequent IgE production^29,30^. This raised the possibility that decreased EoE histopathology in *Crlf2^-/-^* mice was due to altered skin sensitization and not due to an effector role for TSLP. In support of this, TSLP was highly induced in the skin during the sensitization phase (Figure 1E) and skin-sensitized *Crlf2^-/-^* mice displayed decreased IgE levels in comparison with skin sensitized WT mice (Figure S3). To determine whether TSLP contributes to the pathogenesis of EoE as an effector cytokine independent of possible alterations in skin sensitization, WT mice were treated with an anti-TSLP neutralizing antibody (M702)^31^ after skin sensitization (Figure 4A). Neutralization of TSLP resulted in decreased accumulation of intraepithelial and lamina propria eosinophils in the esophagus (Figure 4B-C). Neutralization of TSLP significantly decreased basal cell proliferation (Figure 4D-E) and consequent epithelial cell thickening (Figure 4F). Moreover, esophageal lamina propria thickness and vascularization were significantly attenuated by anti-TSLP treatment (Figure 4G-J). Flow cytometric analysis of esophageal cell suspensions obtained from vehicle and oxazolone-challenged anti-TSLP-treated mice confirmed the reduction in eosinophil levels following anti-TSLP treatment and furtherdemonstrated significant decreased esophageal eosinophils and CD4^+^ T cells (Figure 4K-L).

**Figure 4.**
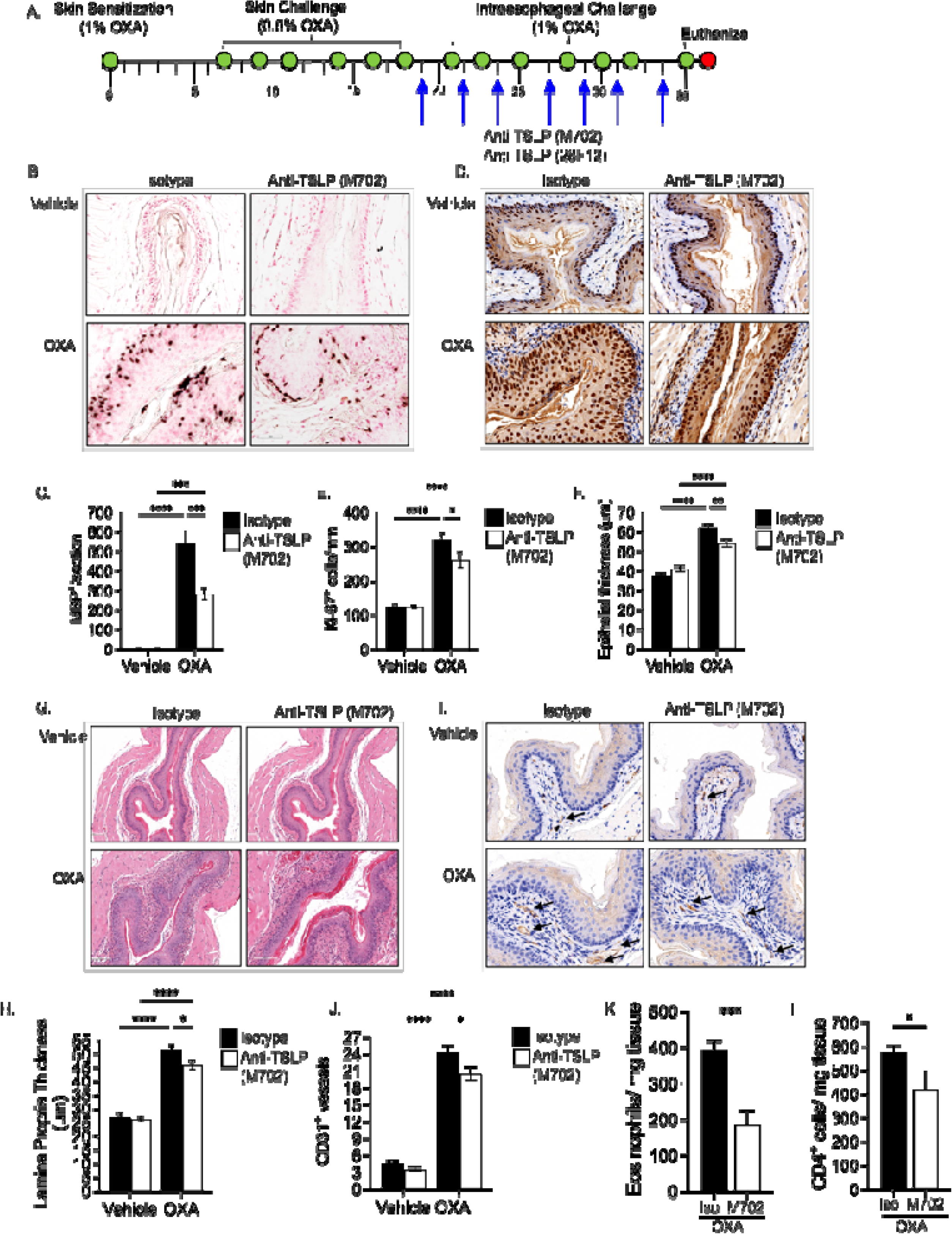
Pharmacological blockade of TSLP regulates eosinophilic infiltration, epithelial cell proliferation and esophageal vascularization in EoE. Experimental EoE was induced in wild type (WT) mice using oxazolone (OXA). Starting on day 19, the mice received two intraperitoneal injections per week of isotype control antibodies or anti-TSLP (clones M702 or clone 28F12, A). On day 36, the mice were euthanized, and esophageal tissues were fixed, paraffin embedded, and slides were generated. The slides were stained with anti-major basic protein (MBP, B) and esophageal eosinophils were quantified (C). Epithelial cell proliferation was determined using anti-Ki67 staining (D, E). H&E-stained slides (G) were analysed for lamina propria (F) as well as epithelial (H) thickness. Esophageal vascularization was determined using anti-CD31 staining (I, J). Single cell suspensions of esophageal tissue were obtained from oxazolone-challenged isotype control or anti-TSLP (M702)-treated mice using enzymatic digestion and the levels of eosinophils and CD4^+^ T cells determined by flow cytometry (K, L). Representative photomicrographs of anti-MBP (B), anti-Ki-67 (D), H&E (G) and anti-CD31 (I) are shown. Data are presented as mean ± SEM and are representative of n=3 experiments conducted with 10-12 mice per group, ns-nonsignificant, *\*-p<0.05, **-p<0.01, ***-p<0.001, ****-p<0.0001*.

To substantiate these findings, TSLP was neutralized using an additional clone of anti-TSLP (BioXcell, clone 28F12). Similar to M702, neutralization of TSLP with a different antibody (clone 28F12) resulted in significant decreases in intraepithelial and lamina propria eosinophils in the esophagus (Figure S4A-B). Furthermore, 28F12-treated mice displayed decreased basal cell proliferation (Figure S4C-D) decreased epithelial cell thickening (Figure S4E) and decreased thickening of the lamina propria (Figure S4F-G).

### RNA sequencing identifies a role for TSLP in IL-13 and IL-1 signaling and epithelial barrier function

Next, global RNA sequencing was conducted on esophageal tissue from vehicle- and oxazolone-challenged mice treated with anti-TSLP antibodies (MC702 or 28F12) or isotype control. Principal component analysis (PCA) identified that most of the transcript variation (77% variance, PC1, Figure 5A) was due to the difference between vehicle-challenged and oxazolone-challenged mice (Figure 5A). Oxazolone-challenged, anti-TSLP-treated mice clustered together, and a 7% variance was observed between oxazolone-challenged anti-TSLP-treated mice and isotype control-treated mice (PC2, Figure 5A).

**Figure 5.**
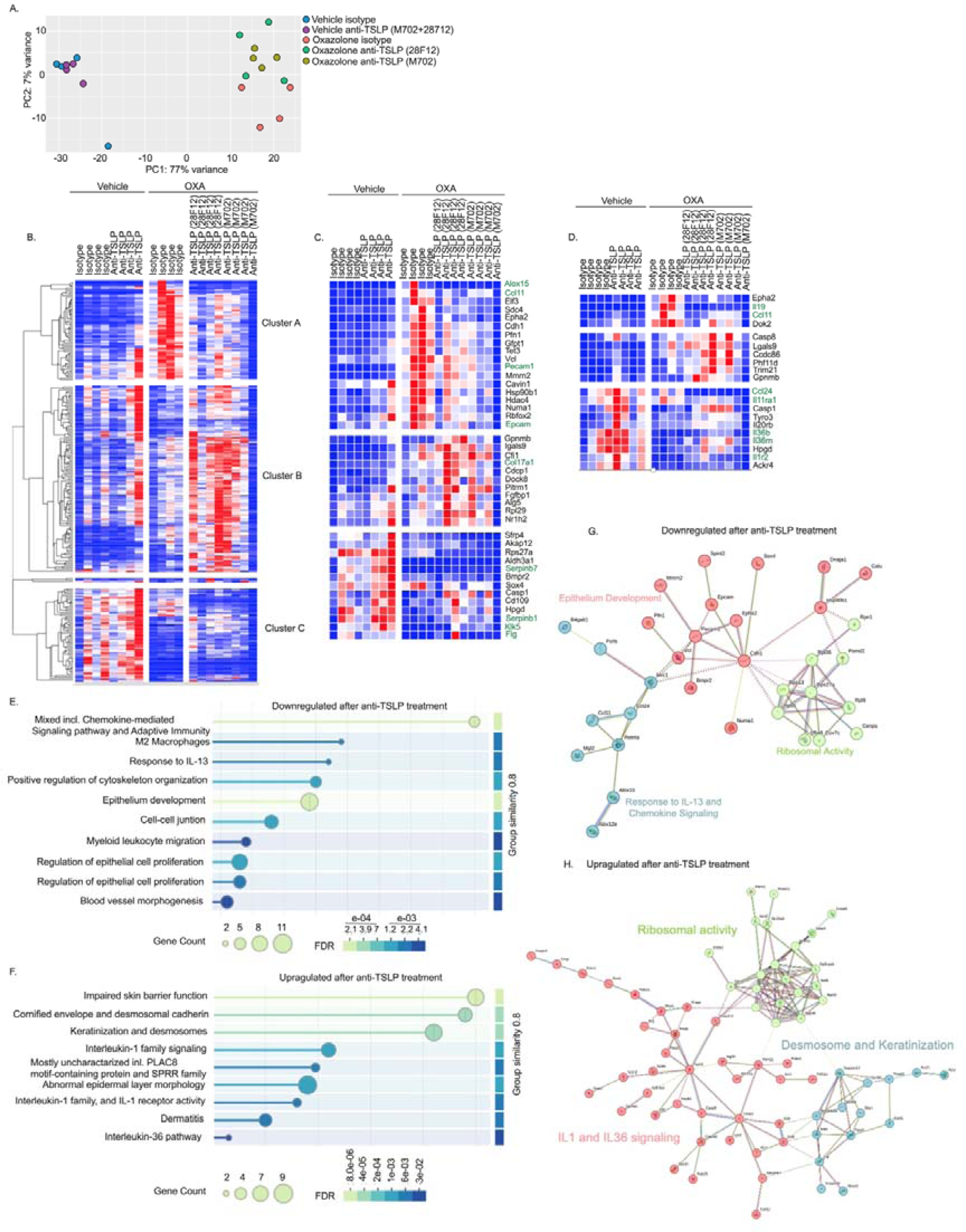
RNA sequencing of anti-TSLP-treated mice. Experimental EoE was induced in wild type (WT) mice. Starting on day 19, the mice received two intraperitoneal injections per week of isotype control antibodies or anti-TSLP (clones M702 or clone 28F12). On day 36, the mice were euthanized, and esophageal tissue was obtained and subjected to bulk RNA sequencing. PCA plot of vehicle- and oxazolone (OXA)-challenged mice treated with either isotype control or anti-TSLP antibodies (clones M702 and/or 28F12) is shown (A). Heat plot analysis of all differentially expressed genes (absolute fold change > 2, p<0.05) (B) is presented. Analysis of the top differentially expressed secreted factors, cytokine receptors, adhesion molecules and extracellular matrix components is shown (C-D). Gene Set Enrichment Analysis (GSEA) and STRING analysis were performed on the DEGs, and comparison of OXA-challenged vehicle-treated mice vs. OXA-challenged anti-TSLP treated mice was performed (E-H). The top up and down regulated pathways, which were regulated by TSLP are presented (E-F). In C, D and E, each lane represents a different mouse.

Comparison between oxazolone-challenged anti-TSLP-treated mice and oxazolone-challenged isotype-control treated mice identified that treatment with the M702 antibody regulated the expression of 195 genes (135 upregulated and 60 downregulated, Table S1) whereas treatment with 28F12 regulated the expression of 149 genes (99 upregulated and 50 downregulated, Table S2). Heat plot representation of the DEGs identified 3 distinct clusters, which were shared between M702- and 28F12-treated mice (Figure 5B). These clusters include: 1) DEGs that were upregulated in isotype-treated OXA and downregulated following anti-TSLP treatment (Cluster A); 2) DEGs that were unchanged in isotype-treated OXA and upregulated following anti-TSLP treatment (Cluster B); 3) DEGs that were downregulated in isotype-treated OXA and relatively upregulated following anti-TSLP treatment (Cluster C).

Anti-TSLP treatment (either with M702 or 28F12) regulated the expression of various cytokines and chemokines and/or their receptors, that have been previously associated with EoE. These include molecules involved in eosinophil migration and accumulation (*Ccl11, Ccll24*) and esophageal epithelial differentiation (*Il36rn, Il36b, Il1r2*)^32^ (Figure 5C). In addition, anti-TSLP treatment downregulated the expression of *Alox12e, Alox15, Il19* and *Mgl2*, which are increased in human EoE and are associated with macrophages and mast cells ^33,27,34^ (Figure 5C-D).

Bioinformatics analysis demonstrated that anti-TSLP treatment downregulated IL-13 signaling associated pathways, M2 macrophages and regulation of migration (Figure 5E-F) and restored gene expression of pathways associated with barrier function, keratinization and desmosome formation, IL-1 receptor activity (Figure 5G-H).

To partially validate the transcriptomics data, key proteins from pathways predicted to be regulated by TSLP (e.g., chemokines) were measured using a protein cytokine array. Consistent with our bioinformatics analysis, anti-TSLP treatment decreased the expression of various chemokines including CCL9, CCL11, CCL22 (Figure S5). Moreover, expression of soluble (s)TNFR1 and sTNFR2 as well as granzyme B were reduced by anti-TSLP treatment. Levels of sIL-2Rα, which has been proposed as a diagnostic market for hyper eosinophilic diseases^35^, were also decreased by anti-TSLP treatment (Figure S5).

### TSLP stimulates IL-13 production in mucosal mast cells

The relatively high expression of CRLF2 on the surface of esophageal MCs (Figure 2) and increased MC levels in human EoE^36–39^, suggest a role for TSLP activation of MCs in EoE. Notably, *Mcpt1* and *Mcpt2*, two highly expressed transcripts in mouse esophageal MCs^40^, were among the top 3 proteases (of 59) that were induced in the esophagus following challenge with oxazolone (Figure 6A). Flow cytometric analysis of single-cell suspensions from enzymatically digested saline- and oxazolone-challenged mice revealed increased MC levels in experimental EoE (Figure 6B). The experimental EoE model used in these studies is critically dependent on IL-13 signaling via IL-13Rα1^25^. The finding that *Crlf2^-/-^* mice and anti-TSLP-treated mice were protected from EoE suggested a role for TSLP as an upstream regulator of IL-13 production. Since esophageal MCs are a predominant cellular source for IL-13^4^, we assessed whether TSLP could stimulate IL-13 production in mouse bone marrow-derived MCs. To this end, MCs were grown under conditions that promoted a mucosal MC phenotype^41^, which includes the induction of *Mcpt1* and *Mcpt2*. Activation of mucosal MCs with TSLP induced the secretion of IL-13 (Figure 6C). TSLP did not promote IL-13 expression from BM-derived MCs that were grown to display a connective tissue-like phenotype (Figure 6D).

**Figure 6.**
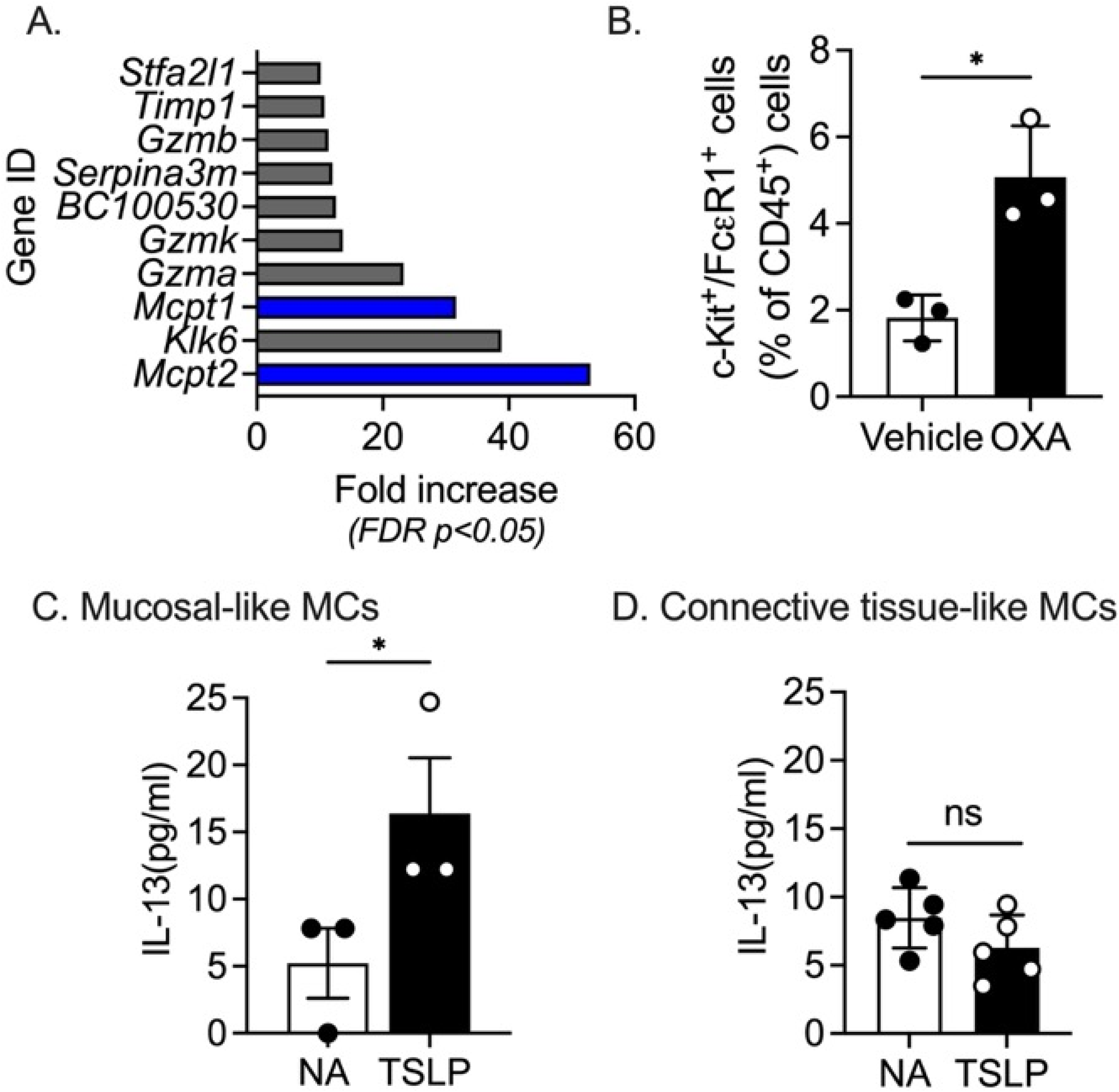
Experimental EoE is characterized by a mucosal mast cell that secrete IL-13 upon stimulation with TSLP. WT mice were treated with vehicle or oxazolone (OXA) to induce experimental EoE. On day 36, the mice were euthanized, and esophageal tissue was obtained and subjected to bulk RNA sequencing. Analysis of the top ten upregulated proteases is shown (p < 0.02, >2-fold, A). Single-cell suspensions were prepared from enzymatically digested esophageal samples and stained with DAPI, anti-CD45, anti– c-Kit, and anti-FcεR1. Thereafter, the percentage of c-Kit^+^/FcεR1^+^ cells were determined (B). Bone marrow-derived mast cells (MCs) were grown and differentiated into a mucosal or connective tissue phenotype (C-D). Thereafter, the MCs were stimulated with TSLP (50 ng/mL) and the secretion of IL-13 was determined by ELISA (C-D). Data are from n = 3. Data are representative of n = 3 conducted in 3-5 technical repeats, ns-nonsignificant, *-*p < 0.05*.

## Discussion

Emerging clinical and experimental evidence emphasize that epithelial cell-driven alarmins including IL-33 and TSLP are at the center of EoE pathogenesis^2,42^. Nonetheless, the contribution of each alarmin and the potential responding cellular compartment involved in the development of EoE is still unclear. Herein, we demonstrate that experimental and human EoE are characterized by increased expression of IL-33 and TSLP. Furthermore, we demonstrate that esophageal MCs highly express TSLP and IL-33 receptors. Pharmacological blockade of TSLP in experimental EoE resulted in decreased esophageal basal cell hyperproliferation, eosinophilia, edema and vascularization. Ablation of IL-33 in experimental EoE resulted only in decreased infiltration of eosinophil to the esophagus, suggesting non-redundant functions for TSLP and IL-33 in EoE. RNA sequencing analyses of esophageal tissues that were treated with two distinct anti-TSLP antibodies identified a role for TSLP in IL-13 signaling, IL-1-dependent pathways, epithelial barrier integrity and chemokine expression. Mechanistically, in vitro activation of mucosal MCs with TSLP resulted in increased IL-13. These data suggest unique roles for TSLP and IL-33 in EoE that may be clinically translated for therapeutic targeting of TSLP in EoE.

TSLP is considered an early upstream regulator of various type 2 immune responses. Upon breach of the mucosal barrier^6^, TSLP is rapidly expressed and released by the activated epithelium and causally contributes to the pathogenesis of asthma, allergic rhinitis, and EoE^6,43^. Indeed, genome-wide association studies (GWAS) have repeatedly determined that genetic susceptibility for EoE is linked to genetic variants in the *TSLP* gene at 5q22^7^; in fact, this is the strongest region of EoE genetic susceptibility across the whole human genome^44^. Directly related, the human *TSLP* locus 5q22 represents a distinguished risk allele for allergic diseases^45^, as determined by multiple, independent GWAS^46,47^. Yet, the linkage of *TSLP* with EoE occurs at an independent location in the genome compared with the risk of other allergic diseases and is present even when controlling for the allergy trait of EoE^48^. Moreover, children carrying genetic variants in *TSLP* and *IL4* display increased risk for EoE compared with those with a risk allele in only one of these genes^49^. The critical roles of TSLP in allergic diseases are best exemplified by interventional clinical trials evaluating inhibition of TSLP in asthma with tezepelumb, an anti-TSLP neutralizing antibody. For example, attenuation of early and late asthmatic responses was observed in response to inhaled allergen^50^, and reduction in annual exacerbation rates was observed in multiple studies^51^, leading to approval of Tezepelumab by the US FDA for the treatment of severe asthma^52^. Thus, defining the roles of TSLP in EoE may provide timely rationale for utilizing anti-TSLP therapeutics in EoE. Our data show that TSLP and IL-33 expression are differently regulated in distinct organs. The expression of TSLP in the esophagus is found under baseline conditions and is increased at later stages of the disease following repetitive intraesophageal challenges with oxazolone. IL-33 is rapidly induced after oxazolone challenge. Interestingly, in the skin, TSLP is induced following allergic sensitization with oxazolone whereas IL-33 is not. This unique expression pattern is likely due to differential regulation within specific cell types. While IL-33 is induced and released by cell damage and death, whereas TSLP expression, especially in epithelial cells, can also be upregulated by pro-inflammatory cytokines such as TNF-a and IL-1b^53^.

Despite the finding that TSLP and IL-33 are co-upregulated in experimental EoE, IL-33 expression was TSLP-dependent. IL-33 regulated esophageal eosinophilia but had no role in epithelial cell proliferation, edema and vascularization. In contrast, TSLP promoted esophageal epithelial cell proliferation, eosinophilia, edema and vascularization. This finding is of specific interest since recent data demonstrated that transgenic overexpression of IL-33 in esophageal epithelial cells induced an EoE-like disease^24^ and that intraperitoneal injection of IL-33 induced IL-13-dependent esophageal eosinophilia and epithelial cell proliferation^22^. Our data suggest that induction of IL-33 may be sufficient but not required to induce EoE under certain circumstances. Notably, previous data demonstrated that IL-33-induced EoE is independent of eosinophils^26^. Similarly, oxazolone-induced EoE was also independent of eosinophils. Thus, the roles of TSLP in this model were also eosinophil independent. These findings corroborate data from recent clinical trials targeting eosinophils in EoE showing a lack of effect on clinical symptoms^54^, and support the clinical relevance of these experimental models. Furthermore, the results suggest that different EoE patients may respond differently to anti-TSLP or anti-IL-33 treatment despite the presence and/or upregulation of both alarmins in the esophagus.

IL-13 is a key effector cytokine that is involved in EoE pathogenesis; this was demonstrated in multiple in vivo studies and in human patients treated with therapeutics targeting IL-4Rα, which blocks IL-13 signaling via the type 2 IL-4R, or treated with therapeutics blocking IL-13^55,56^. Importantly, the histopathology (i.e., epithelial cell proliferation, edema, vascularization, eosinophilia), which is observed in oxazolone-induced EoE, is entirely dependent on IL-13 signaling via IL-13Rα1^25^. Thus, it is likely to assume that neutralization of TSLP resulted in decreased expression of IL-13 and subsequent reduced IL-13-driven activities. This notion was further supported by RNA sequencing analysis demonstrating decreased expression of various pathways that converge on IL-13 and epithelial cell “gene signatures”. Anti-TSLP treatment decreased the expression of various known IL-13-regulated genes including *Ccl11, Ccl24, Mgl2, Retnla, Alox15*. Furthermore, anti-TSLP treatment increased the expression of IL-13-regulated barrier function-associated genes such as *Flg* and *Klk5*^3,57^. Our study bears the limitation that we could not demonstrate that anti-TSLP reduced IL-13 protein expression in the esophagus. This is a general limitation of the model, since IL-13 protein is not detected in esophageal lysates even after challenge with oxazolone (data not shown). Nonetheless, the critical involvement of IL-13 in this experimental model was previously proven by three independent methods: 1) Failure to induce experimental EoE in *Il13ra1^-/-^* mice; 2) therapeutic neutralization of IL-13 in experimental EoE; and 3) Induction of experimental EoE in mice that lack *Il13ra1* specifically in esophageal (and skin) epithelial cells^25^.

TSLP has been shown to activate multiple cells that can promote type 2 immune responses. It can directly upregulate co-stimulatory molecules on dendritic cells, which are required for the polarization of T cells into T helper 2 (Th2) cells^11,12^. TSLP can also increase the proliferation of Th2 cells and enhance the release of Th2 cell-associated cytokines and chemokines from MCs, innate lymphoid cells (ILCs), and macrophages^13–17^. Our data suggests that TSLP-activated MCs promote EoE, at least in part by secretion of IL-13. Analysis of RNA sequencing data of the esophagus following induction of experimental EoE identified a striking elevation of esophageal *Mcpt1 and Mcpt2* expression, which were associated with increased levels of esophageal MCs. These data are consistent with increasing evidence suggesting a role for MCs in EoE^36^. Patients with active EoE displayed increased numbers of esophageal MCs and increased degranulation^36^, which correlated with key disease symptoms, such as dysphagia^39^. Elevated esophageal expression of MC CPA3 and chymase associated with a steroid-resistant subtype of EoE^4^, whereas reduced MC numbers correlated with a positive clinical response to emerging biological treatments^58,59^. Mechanistically, MCs have a broad impact on the pathophysiology in EoE including disruption of the esophageal epithelial barrier^38^, odynophagia^37^, and dysphagia through increased esophageal smooth muscle contraction^39^. Furthermore, a pathogenic role for MCs has been reported in an aeroallergen-induced EoE model^60^. Recently, a heterogeneous group of esophageal MCs were identified in EoE patients^4^. Among these MC populations, a transient, proinflammatory, activated subset, marked as *Kit^hi^/Il1rl1^hi^/Fcer1^low^,* was detected^4^. This MC subset was a prominent source of esophageal IL-13 mRNA and protein^61,62^. Our data show that esophageal MCs highly express TSLPR (in human and mice) and that activation of mucosal MCs results in secretion of IL-13. Moreover, anti-TSLP treatment decreased the expression of various genes that were shown to be expressed in MCs (and macrophages) especially in the context of allergy, including *Retnla, Il19, Mgl2* and *Alox15*. Future studies aimed to determine the contribution of MC-driven IL-13 in EoE should be conducted.

Collectively, these data demonstrate distinct roles for TSLP and IL-33 in EoE and suggest that pharmacological targeting of TSLP may serve as an opportunity for the treatment of EoE and perhaps additional allergic diseases.

## Acknowledgments

We wish to thank all the members of the Munitz lab for their constructive comments.

## Author Contributions

These authors contributed as follows:

AD, JDS and AM-conception and/or design of the work

AD, AW, SA, MI, NR-Data Collection

AD, AW, SFZ, JRP, KSG, JDS, IZB, MER and AM-Data analysis and interpretation.

AD, SZ, JDS, JRP, KSG, MER and AM-Drafting the article

AD, AW, SA, MI, NR, JRP, KSG, JDS, IZB, SFZ, MER and AM - Critical revision of the article

AM-Final approval of the version to be published

