## Supplemental Data for "Distinct roles for thymic stromal lymphopoietin (TSLP) and IL-33 in experimental eosinophilic esophagitis"

**Short Title:** TSLP in eosinophilic esophagitis

**Key words:** Allergy, Eosinophilic Esophagitis, TSLP, IL-33

[*]**Corresponding author:** Ariel Munitz, PhD, Department of Clinical Microbiology and Immunology, The Sackler School of Medicine, Tel Aviv University, Ramat Aviv 69978, Israel. Tel. (Office): +972-3-640-7636, Fax: +972-3-640-9160,; ORCID: https://orcid.org/0000-0003-1626-3019;

**Materials and methods:**

**Mice**

Wild-type (WT) C57BL/6 mice were obtained from Harlan Laboratories (Rehovot, Israel) and grown in-house. *Il33*$^{-/-}$ mice[1], were obtained from Dr Jakub Abramson (Weizmann Institute of Science, Rehovot, Israel). *Crlf2*$^{-/-}$ mice were All experiments were reviewed and approved by the Animal Care Committee of Tel Aviv University and were performed in accordance with its regulations and guidelines regarding the care and use of animals for experimental procedures. In all experiments, age-, weight-, and sex-matched mice were used and housed under specific-pathogen-free conditions according to protocols approved by the Tel-Aviv University Institutional Animal Care Unit.

**Experimental eosinophilic esophagitis (EoE)**

Experimental EoE was induced as previously described[2]. Briefly, mice (male c57BL6 mice, 6–8 weeks) were skin sensitized with 15 µL of 1% Oxazolone (Sigma, #E0753) solution dissolved in acetone on both ear flanks (60 µL per mouse). After six additional skin challenges (0.5% OXA in acetone), the mice were bled and serum IgE was quantitated by ELISA (BD Bioscience, #555248). On Day 18, the mice were intraesophageally challenged (7 challenges) with 200 µL OXA (1% in a 1:2 ratio of olive oil and 95% alcohol, respectively) using a plastic feeding tube that reaches the mid-esophagus (INSTECH, #ITH-FTP-22-25-50, 22ga X 25 mm) and has been modified for intraesophageal administration by puncturing 8 holes with a 14G needle. On days 29-36 the mice were euthanized.

**In vivo neutralization experiments**

On Day 19 of the experimental EoE protocol, mice were intraperitoneally injected with 150 µg of anti-mouse TSLP neutralizing antibody 28F12 (BioXcell), anti-mouse TSLP neutralizing antibody M702 (Amgen)[3], anti-mouse IL-5 (Clone TRFK4, BIoXcell)[4],150 µg isotype control rat IgG2a (BioXcell, #BE0089), or 150 µg isotype control rat IgG1 (BioXcell,# BE0088). Additional injections were given twice a week until day 36. On day 36 the mice were euthanized, and samples were taken for analysis.

**Histology and immunohistochemistry**

Esophageal tissue was harvested, fixed with 4% formaldehyde (Bio-lab ltd. Israel), and paraffin-embedded. Thereafter, slides of paraffin-embedded sections (5 µm) were processed and stained with H&E (Sigma, Israel), anti-MBP (kindly provided by Dr. Elizabeth A. Jacobsen, Mayo Clinic, Scottsdale, AZ), anti-Ki-67 (Novus, # NB110-89717SS), anti-mouse-TSLP (Merc, #ABT330) and anti-mouse IL-33 (Abcam, #EPR17831). Thereafter, the slides were scanned using the Aperio slide scanner (Leica) and analyzed using Image Scope (Leica) and ImageJ.

**TSLP and IL-33 ELISA**

Esophageal tissues were harvested and normalized by weight. Total protein was extracted using Pierce IP Lysis Buffer (Thermo Scientific, #87787, USA). Tissue lysates were concentrated using 10K Centrifugal Filter Units (Merck, # UFC501024 Cairo, Egypt) and the presence of TSLP or IL-33 was determined by ELISA. TSLP was detected using mouse TSLP duoset ELISA (R&D Systems, # DY555). IL-33 was detected by mouse IL-33 duoset ELISA (R&D Systems, # DY3626). All ELISA assays were performed according to manufacturer manuals.

**Enzymatic digestion of esophageal biopsies**

A single section from the distal esophagus was collected into RPMI-1640 medium (Thermo, # 11875093) supplemented with 10% FBS on ice. The biopsy was then transferred into EDTA buffer (5mM EDTA, 25mM HEPES, 10% FBS in HBSS) for 15 minutes at 37°C, washed once with PBS, minced, and then subjected to collagenase A (2.4 mg/mL, Roche, # 10103586001) digestion for 30 minutes at 37°C. The resulting suspension was diluted with ice-cold PBS, passed through a 19-gauge needle, filtered through a 70μm cell strainer, and washed with ice-cold PBS.

**Flow cytometry**

Single-cell suspensions were prepared from enzymatically digested esophageal specimens. Cells were stained with a panel of fluorochrome-conjugated antibodies targeting specific immune cell markers. The panel included CD45-PE/Cy7 (Clone: 30-F11), CD11b-PerCP/Cy5.5 (Clone: M1/70), CD45-APC-eFluor780 (Clone: 30-F11) (eBioscience, San Diego, CA, USA); SiglecF-PE (Clone: E50-2440) (BD Biosciences, San Jose, CA, USA); Ly6C-PE/Cy7 (Clone: HK1.4), F4/80-AF700 (Clone: BM8), CD3ε-PE/Cy7 (Clone: 145-2C11), CD4-PerCP/Cy5.5 (Clone: GK1.5), T cell receptor γ/δ–APC (Clone: GL3), CD8α-BV510 (Clone: 53-6.7), TSLPR-APC, ST2-APC, and Rat IgG 2a-APC (BioLegend, San Diego, CA, USA); and Ly6G-APC/Cy7 (Clone: 1A8) (Bio Gems, Westlake Village, CA, USA). Cell viability was assessed using 4′,6-diamidino-2-phenylindole (DAPI) (Sigma-Aldrich, Rehovot, Israel). Absolute cell counts were determined with Flow-Count Fluorospheres (Beckman Coulter, Brea, CA, USA) following the manufacturer’s protocol. Data acquisition was performed on a Gallios flow cytometer (Beckman Coulter, Brea, CA, USA), ensuring appropriate voltage and compensation settings. A minimum of 10,000 live events were collected for each sample. Flow cytometric data were analyzed using Kaluza software (Beckman Coulter, Brea, CA, USA).

**Bulk RNA sequencing**

The esophagus was dissected, and 1 cm pieces from segments 2 and 3 were used for RNA extraction using TRIzol reagent (Invitrogen, #15596026). RNA was quantified using a Qubit Flex Fluorometer (Invitrogen) with the Equalbit RNA High Sensitivity Assay Kit (Vazyme, #EQ211). CEL-Seq2 libraries were prepared as described[5], with modifications to use 2 ng of purified RNA as input instead of single cells. Sequencing was performed on an Illumina NextSeq2000 platform using P2 100 cycles (Read1-12; Index1-6; Index2-0; Read2-65). Reads were demultiplexed following the CEL-Seq2 pipeline with parameters: min_bc_quality = 10, bc_length = 6, umi_length = 6, and cut_length = 70. Quality control was performed using FastQC (v0.11.9), and reads were trimmed for adapters and poly-A tails using CUTADAPT (v4.4) with a minimum Phred score of 20 and a minimum length of 25 bp. Reads were aligned to the Mus musculus GRCm39 genome using HISAT2 (v2.2.1) with up to two mismatches allowed per read, and only uniquely mapped reads were retained using SAMtools (v1.13). Gene-level read counts were obtained using HTSeq (v2.0.4) in 'union' mode, and normalization and differential expression analysis were conducted using DESeq2 (v1.28.0). Sample preparation and sequencing were conducted by the "Technion Genome Center”, Life Science and Engineering Interdisciplinary Research Center, Technion, Haifa, Israel.

**Bioinformatics analysis**

The differential expression (DE) of the sequenced genes was calculated using the DEseq2 package in R language on RStudio. Genes were regarded as differentially expressed if they presented a false discovery rate (FDR) adjusted P-value below 0.05, and $log2(fold\ change) \pm 0.58$. The samples were presented using principal component analysis (PCA) created with the ggplot R package. Volcano plots (that visualize results of differential expression analyses) were created using the Enhanced Volcano package in R.

Expression heat maps of selected genes of interest (GOIs) were generated using the Morpheus software (Broad Institute). The expression data was subsetted from the sequencing dataset and scaled per gene (row) to enable direct comparison across genes. Hierarchical clustering was performed using the built-in clustering method in Morpheus. The heatmaps visually represent the relative expression levels of GOIs across experimental conditions or samples.

To explore protein-protein interactions (PPIs) of selected GOIs, analyses were conducted using the STRING database. Genes were queried to identify known and predicted interactions based on experimental evidence, computational predictions, and curated databases. The STRING output was used to construct interaction networks, aiding the identification of functional pathways and complexes potentially involved in the observed experimental outcomes.

**Bone marrow-derived mucosal and connective tissue mast cell (MC) culture and activation**

Bone marrow cells were harvested from both femoral bones of a mouse in a sterile environment. The bones were flushed with RPMI-1640 medium supplemented with 10% FBS, 100 U/mL penicillin, 100 μg/mL streptomycin, 2 mM L-glutamine, 1x nonessential amino acids, 10 mM HEPES, 50 μM 2-Mercaptoethanol, 20 ng/mL murine recombinant SCF (Thermo, #250-03) and 20 ng/mL murine recombinant IL-3 (Thermo, #213-13). To differentiate MCs into a mucosal phenotype, the media was supplemented with additional 20 ng/mL murine recombinant IL-9 (Thermo, #219-19), and 2 ng/mL TGF-β (Thermo, #100-21)[6]. To differentiate MCs into a connective tissue phenotype the media was supplemented with 5 ng/mL IL-4 (Thermo, Gibco™ 2141420UG)[6]. The cell suspension was collected in a 50 mL tube and passed through a 70 μm cell strainer to remove bone fragments and debris. The cell suspension was plated in 10 mL of the complete RPMI-1640 medium in a 100 mm Petri dish. Cells were cultured at 37°C in a humidified atmosphere containing 5% $CO_2$. On days 1, 3, 5, 7, 10, 14, 18, 22, and 26, cells were spun down, re-suspended in fresh medium, and transferred to new culture dishes to refresh the cells and discard any adherent cells. Starting from day 7, cells were transferred to larger culture flasks (252 mm), and the cell concentration was adjusted to 1.5 × $10^6$ cells/mL. The culture was maintained for 21-28 days. At the end of the culture period, cell purity was assessed by flow cytometry, targeting FcεRI and c-Kit expression, and was expected to achieve a purity of 95-98%.

On day 21, the cells were harvested, counted, and re-suspended at 1 × $10^6$ cells/mL concentration in a complete RPMI-1640 medium. The cells were then seeded into a 24-well plate at a density of 0.2 million cells per well. Each well was stimulated with thymic stromal lymphopoietin (eBioscience, #14-8498-62) at a final concentration of 50 ng/mL. The plates were incubated at 37°C in a humidified atmosphere containing 5% $CO_2$ for 24 hours. After the 24-hour stimulation period, culture supernatants were collected and centrifuged at 300 × g for 10 minutes to remove cellular debris. The levels of IL-13 in the supernatants were measured using an anti-mouse IL-13 ELISA kit (DY413, R&D), following the manufacturer's instructions.

**Protein Array**

Following the manufacturer's protocol, protein expression was analyzed using the Abcam Mouse 97 Cytokine Antibody Array (ab169820). Esophageal tissues from four groups (vehicle + isotype antibody, vehicle + anti-TSLP antibody, OXA + isotype antibody, and OXA + anti-TSLP antibody) were homogenized in Pierce™ IP Lysis Buffer supplemented with protease and phosphatase inhibitors. Lysates were centrifuged at 14,000 × g for 10 minutes at 4°C, and protein concentrations were determined using a BCA Protein Assay Kit. Equal amounts of protein from each sample were applied to pre-blocked membranes and incubated overnight at 4°C. Following washing, biotinylated secondary antibodies and HRP-streptavidin were applied sequentially, and chemiluminescent signals were visualized using ECL reagents and the ChemiDoc Imaging System. Signal intensities were normalized to internal controls for analysis.

**Statistical analysis**

*P* values of mouse data sets were determined by one-way analysis of variance (ANOVA), unpaired two-tailed Student's *t-test* with a 95% confidence interval. All statistical tests were performed with GraphPad Prism V10 software. Data are shown as mean ± SEM. *-p < 0.05; **-*p* < 0.01; ***-*p* < 0.001.

#### Figure legends

##### Figure S1. Gating strategy of mouse esophageal cell

Single cell suspensions were obtained from the esophagus of vehicle and oxazolone treated wild type mice and stained with the indicated antibodies and viability dye. The gating strategy for identification of Monocytes (Mono), Neutrophils (Neut), macrophages (Mac), dendritic cells (DC), basophils (Bas), mast cells (MCs), eosinophils (Eos), epithelial cells (Epi), endothelial cells (Endo) and fibroblasts (Fib) is shown (A). Histogram plot analysis depicting the expression of ST2 (A) and CRLF2 (B) on the surface of esophageal MCs is shown. Data are representative of one esophagus from at least n=3 different mice.

##### Figure S2. Eosinophils are dispensable for induction of EoE

Experimental EoE was induced in wild type (WT) mice using oxazolone (OXA). Starting on day 19, the mice received two intraperitoneal injections per week of isotype control antibodies or anti-IL-5 (clone TRFK, A). On day 36, the mice were euthanized, and esophageal tissues were fixed, paraffin embedded, and slides were generated. The slides were stained with anti-major basic protein (MBP) and esophageal eosinophils were quantified (B). Epithelial cell proliferation was determined using anti-Ki67 staining (C). H&E-stained slides were analysed for epithelial and lamina propria thickness (D, E). Data are presented as mean ± SEM and are representative of n=2 experiments conducted with 10-12 mice per group, ns-nonsignificant, *- *p<0.05*, **-*p<0.01*, ***-*p<0.001*, ****- *p<0.0001.*

##### Figure S3. Decreased serum IgE in skin sensitized *Crlf2*$^{-/-}$ mice

Wild type (WT) and *Crlf2*$^{-/-}$ mice were skin sensitized using oxazolone (OXA). On day 17, the mice were bled and levels of serum total IgE were determined by ELISA. Data are presented as mean ± SEM and are representative of n=3 experiments conducted with 10-12 mice per group, ns-nonsignificant, *- $p<0.05$.

**Figure S4. Pharmacological blockade of TSLP regulates eosinophilic infiltration, epithelial cell proliferation and esophageal vascularization in EoE.**

Experimental EoE was induced in wild type (WT) mice using oxazolone (OXA). Starting on day 19, the mice received two intraperitoneal injections per week of isotype control antibodies or anti-TSLP (clone 28F12). On day 36, the mice were euthanized, and esophageal tissues were fixed, paraffin embedded, and slides were generated. The slides were stained with anti-major basic protein (MBP, A) and esophageal eosinophils were quantified (B). Epithelial cell proliferation was determined using anti-Ki67 staining (C, D). H&E-stained slides (F) were analysed for epithelial and lamina propria thickness (E, G). Data are presented as mean ± SEM and are representative of n = 2 experiments conducted with10-12 mice per group, ns-nonsignificant, *- $p<0.05$, **-$p<0.01$, ***-$p<0.001$, ****- $p<0.0001$.

**Figure S5. Pharmacological blockade of TSLP regulates esophageal chemokine expression**

Experimental EoE was induced in wild type (WT) mice. Starting on day 19, the mice received two intraperitoneal injections per week of isotype control antibodies or anti-TSLP (clone M702). On day 36, esophageal tissues were obtained, and protein lysates were prepared and subjected to antibody-based protein array. Protein arrays were quantitated by subtracting the background expression and proteins that were downregulated by anti-TSLP treatment are shown. Data are from n = 4 arrays for vehicle treated groups (2 for vehicle isotype and 2 for vehicle anti-TSLP) and n = 6 arrays for oxazolone (OXA)-treated groups (4 for vehicle isotype and 4 for vehicle anti-TSLP), *- $p<0.05$, **-$p<0.01$, ***-$p<0.001$, ****- $p<0.0001$.

### Figure S1.

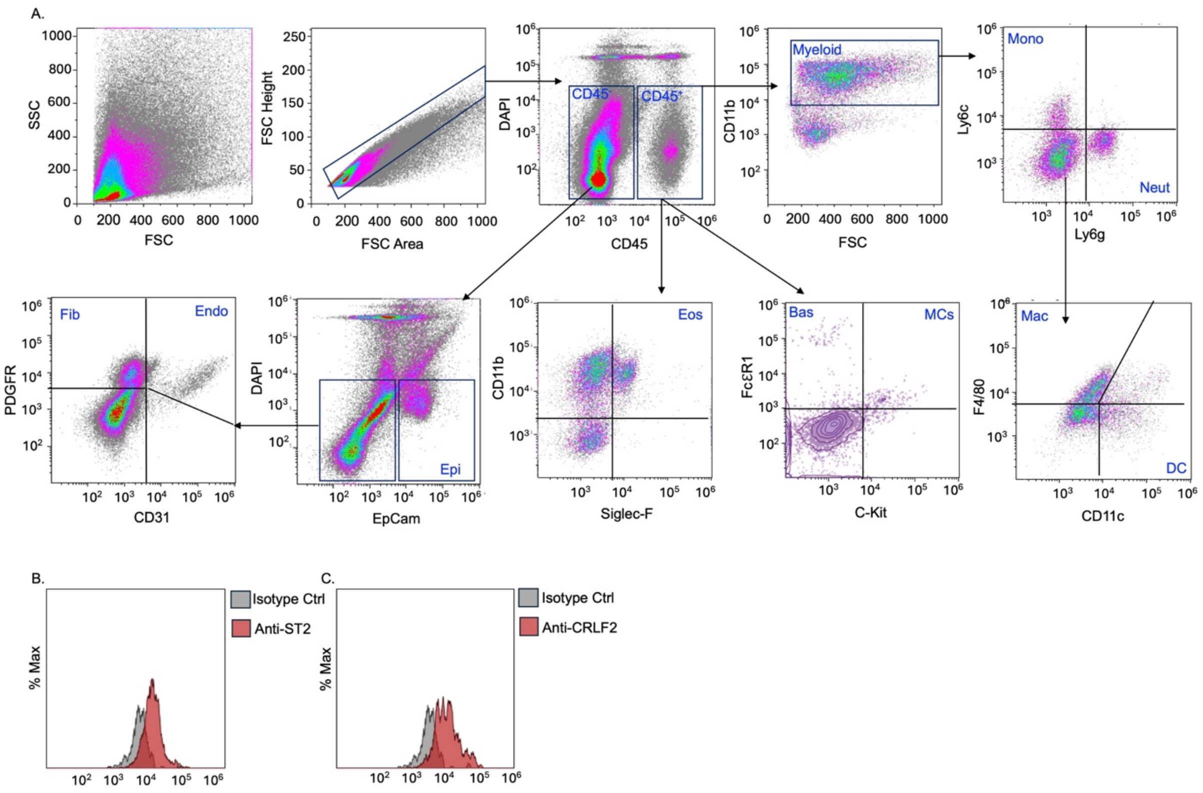
A.
SSC
FSC
FSC Height
FSC Area
DAPI
CD45
CD45⁻
CD45⁺
CD11b
Myeloid
Ly6c
Ly6g
Mono
Neut
PDGFR
CD31
Fib
Endo
EpCam
Epi
Siglec-F
Eos
FcεR1
C-Kit
Bas
MCs
F4/80
CD11c
Mac
DC
B.
% Max
Isotype Ctrl
Anti-ST2
C.
% Max
Isotype Ctrl
Anti-CRLF2

**Figure S2**

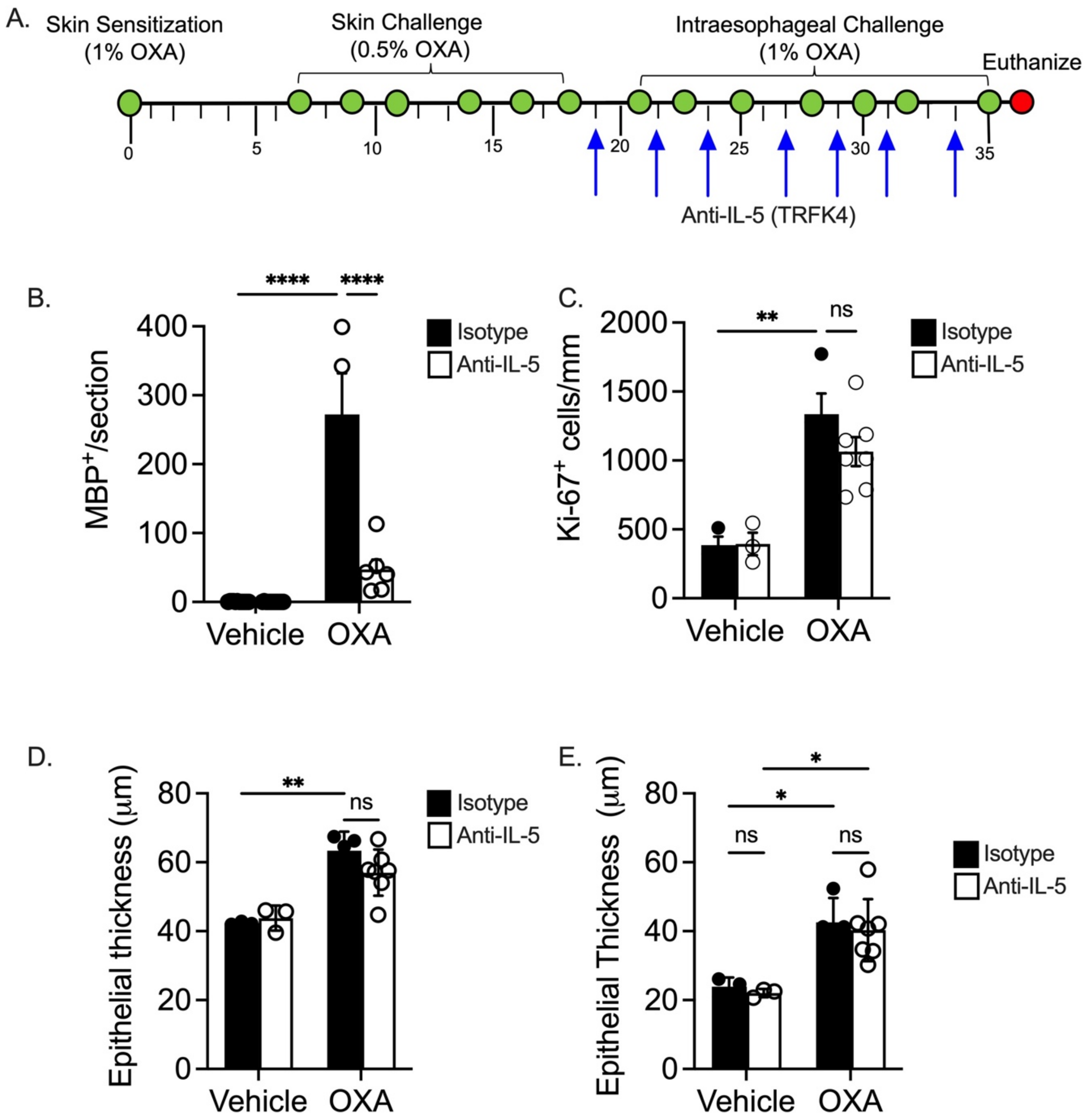
A.
Skin Sensitization
(1% OXA)
Skin Challenge
(0.5% OXA)
Intraesophageal Challenge
(1% OXA)
Euthanize
0
5
10
15
20
25
30
35
Anti-IL-5 (TRFK4)
B.
MBP+/section
****
****
Isotype
Anti-IL-5
Vehicle
OXA
C.
Ki-67+ cells/mm
**
ns
Isotype
Anti-IL-5
Vehicle
OXA
D.
Epithelial thickness (μm)
**
ns
Isotype
Anti-IL-5
Vehicle
OXA
E.
Epithelial Thickness (μm)
*
*
ns
ns
Isotype
Anti-IL-5
Vehicle
OXA

**Figure S3**

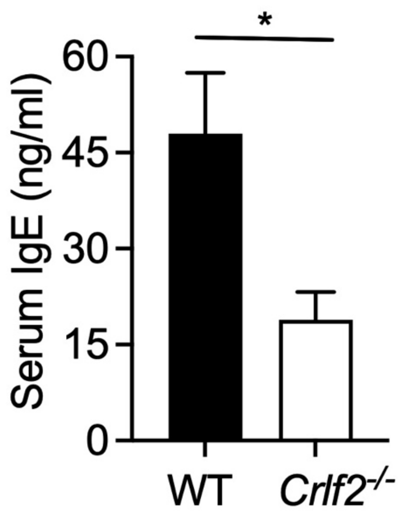

**Figure S4.**

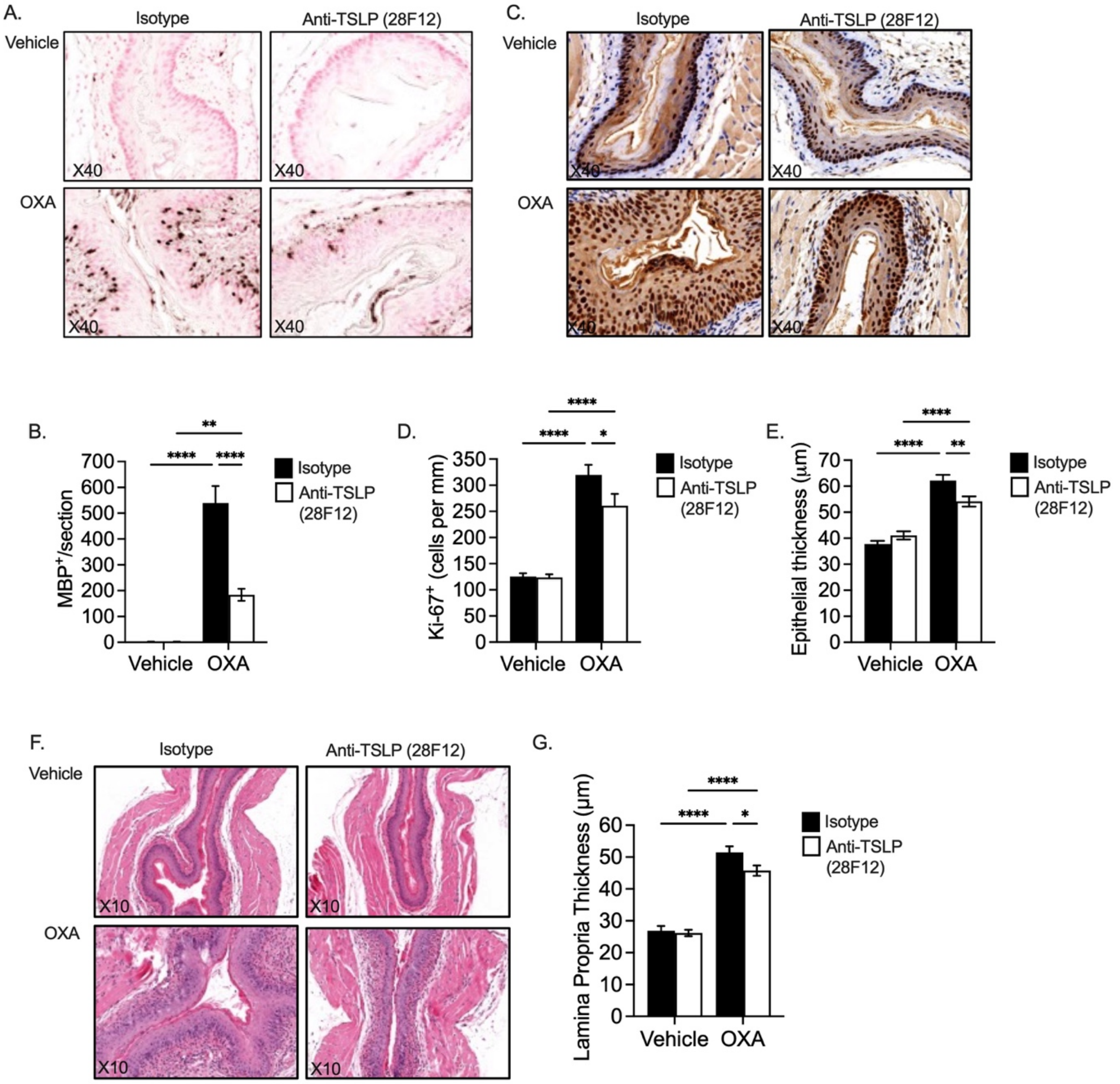

A.
Isotype
Anti-TSLP (28F12)
Vehicle
OXA
X40
C.
Isotype
Anti-TSLP (28F12)
Vehicle
OXA
X40
B.
MBP+/section
Vehicle
OXA
Isotype
Anti-TSLP (28F12)
D.
Ki-67+ (cells per mm)
Vehicle
OXA
Isotype
Anti-TSLP (28F12)
E.
Epithelial thickness (μm)
Vehicle
OXA
Isotype
Anti-TSLP (28F12)
F.
Isotype
Anti-TSLP (28F12)
Vehicle
OXA
X10
G.
Lamina Propria Thickness (μm)
Vehicle
OXA
Isotype
Anti-TSLP (28F12)

**Figure S5**

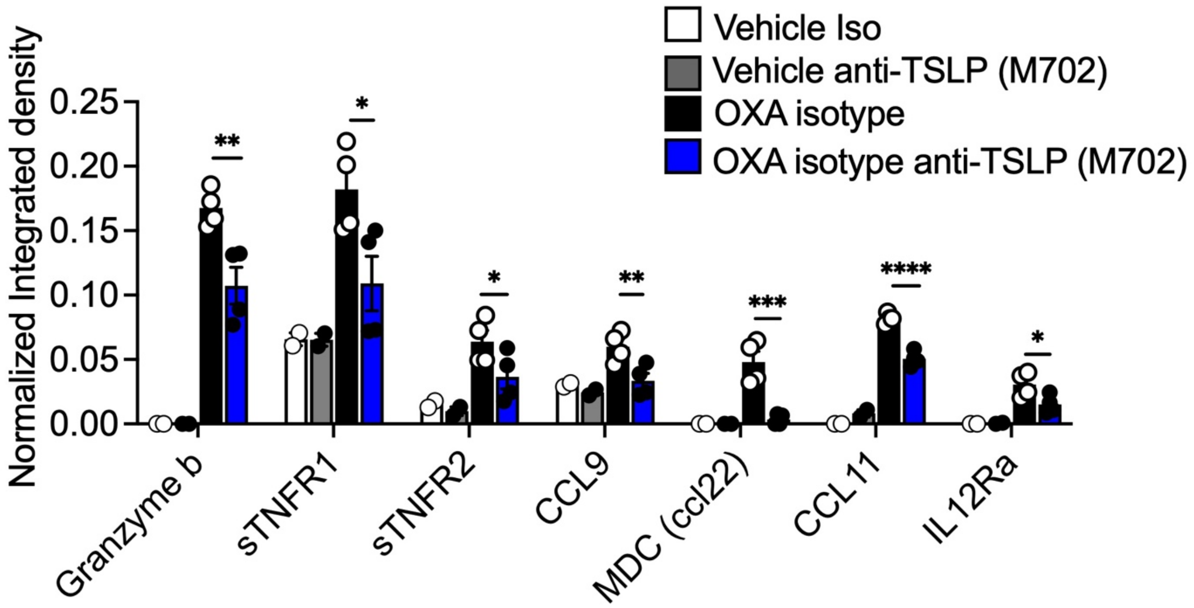

Vehicle Iso
Vehicle anti-TSLP (M702)
OXA isotype
OXA isotype anti-TSLP (M702)
Normalized Integrated density
0.25
0.20
0.15
0.10
0.05
0.00
**
*
*
**
***
****
*
Granzyme b
sTNFR1
sTNFR2
CCL9
MDC (ccl22)
CCL11
IL12Ra
